# Single-cell Morphological Mechanisms-of-Action Profiling using Nanowell Imaging Plates

**DOI:** 10.64898/2026.11.23.768734

**Authors:** Samuel G. Berryman, Kerryn Matthews, Simon P. Duffy, Hongshen Ma

## Abstract

Mechanism-of-action (MoA) analysis is used in drug discovery to prioritize compounds that act through desired pathways while eliminating compounds associated with nonspecific or undesirable activity. Image cytometry provides a promising route for MoA inference by capturing high-dimensional cellular phenotypes at single-cell resolution, but conventional workflows remain limited by unreliable cell segmentation and dependence on human-engineered feature extraction. Here, we present a deep learning-based image cytometry workflow that confines single cells in nanowells to obtain uniformly framed, regularized single-cell images, enabling MoA inference directly from raw images without segmentation or human-engineered features. Using a nanowell-in-microwell platform, we treated cells with 21 chemotherapeutic compounds spanning 11 MoA classes across multiple doses and time points, generating more than 500,000 single-cell images for model training and validation. A ResNet-18 convolutional neural network trained on images annotated by known compound mechanism achieved 85% accuracy in MoA classification. When a second model was trained on compound identity rather than MoA, misclassification patterns frequently linked compounds with shared mechanisms, indicating that the model learned mechanism-level relationships directly from single-cell morphology, without those relationships being specified in advance. Together, these results establish regularized single-cell imaging in nanowells as a scalable and interpretable approach to image-based MoA classification that overcomes key limitations of segmentation-dependent feature analysis.

## Introduction

Mechanism-of-action (MoA) analysis is important in early-stage drug discovery to prioritize compounds that act through desired pathways while eliminating compounds linked to nonspecific, unproductive, or unsafe effects^1,2^. Despite its importance, MoA analysis remains challenging, as many compounds elicit pleiotropic, heterogeneous, and context-dependent cellular responses that are poorly captured by single-target assays^3^. Image cytometry offers a promising alternative because it captures rich, high-dimensional cellular phenotype data at single-cell resolution, which allows compounds with shared biological effects to be grouped by their morphological signatures to provide a scalable framework for MoA inference^4^.

Image cytometry is a microscopy-based approach for quantitative analysis of cellular phenotype at single-cell resolution. The general workflow involves exposing cells to chemical or genetic perturbations, labelling with fluorescent dyes highlighting specific subcellular structures, and imaging using high-content microscopy to capture multidimensional features such as cell size, shape, texture, intensity, and intracellular organization^5,6^. Widely used implementations, such as Cell Painting, employ multiplexed fluorescent dyes label diverse cellular compartments and generate rich morphological readouts across large numbers of perturbations^7,8^. The resulting images are then processed to identify individual cells, extract quantitative descriptors, and assemble high-dimensional phenotypic profiles that can be compared across treatments using dimensionality reduction, clustering, or supervised classification^9,10^. Since compounds that perturb related pathways often induce similar morphological states, image cytometry enables grouping compounds by shared phenotypic signatures to support MoA inference from cellular response rather than target-specific assays alone.

Despite its promise for MoA inference, conventional image cytometry workflows remain constrained by two key analytical bottlenecks: cell segmentation and feature extraction. Accurate segmentation of individual cells is often difficult in practice because of debris, cell overlap, imaging artifacts, poorly defined boundaries, and heterogeneous morphology^11^. These difficulties are exacerbated under the cytotoxic conditions where cell death, debris, and drug-induced morphological changes further degrade segmentation. Segmentation-based workflows are also subject to cell-density bias, as both cell morphology and the reliability of single-cell isolation depend on local cell density, which varies across a conventional well. As a consequence of these challenges, substantial manual tuning and filtering is required to prevent over-segmentation and under-segmentation to obtain usable single cell images for feature extraction^12^. Importantly, errors introduced at this stage propagate directly into downstream analysis by corrupting the measurements assigned to each cell. Even when segmentation is successful, conventional workflows typically rely on predefined, human-engineered descriptors such as size, shape, texture, and fluorescence intensity to represent cellular phenotype. Although these features capture many aspects of morphology, they are limited by assumptions about which image properties are likely to be important and may fail to capture subtle or unexpected phenotypes of true biological importance^13^. Together, the unreliability of segmentation and the incompleteness of human-engineered feature extraction constrain the robustness, scalability, and interpretability of current image-based workflows for MoA inference.

Deep learning approaches are emerging as potential approaches to address key challenges in image cytometry, primarily by automating cell segmentation and feature extraction^14–16^. However, existing feature extraction workflows often retain reliance on predefined or implicitly constrained feature representations, which may overlook phenotypes relevant to compound MoA. An alternative strategy is to use convolutional neural networks (CNNs) to learn feature representations directly from images of compounds with known MoA^17^. However, such learned embeddings introduce a representation bottleneck, compressing complex cellular phenotypes into fixed-length vectors that may discard subtle but biologically relevant signals. More recently, models have been developed to infer MoA directly from microscopy images without explicit feature extraction or segmentation^18,19^. While these approaches avoid segmentation-related errors, they sacrifice single-cell resolution, effectively averaging image signals across populations and limiting the ability to detect subpopulation-specific or partial responses to compound treatment. These limitations highlight the need for a deep learning framework that preserves single-cell resolution through segmentation while eliminating feature extraction, enabling direct learning from whole-cell images without imposing restrictive intermediate representations.

Here, we present a deep learning-based image cytometry workflow that addresses these bottlenecks at the point of image acquisition rather than through downstream computation. By confining single cells within regularly spaced nanowells, we acquire uniformly framed, *regularized* single-cell images in which each cell is presented at a consistent scale, position, and background. This regularized imaging strategy converts unreliable post-hoc segmentation into a controlled acquisition problem and removes a major source of technical variability, enabling direct MoA inference from raw single-cell images without segmentation or hand-engineered feature extraction. Using our previously described nanowell-in-microwell platform^20–23^, single cells are isolated in nanowells, preventing cell overlap and enabling standardized imaging without the need for unreliable cell segmentation. We treated cells with 21 reference chemotherapeutics across multiple doses and time points and acquired more than 500,000 single-cell images to train and validate a ResNet-18 convolutional neural network (CNN) directly on raw images rather than on manually extracted features. A CNN trained on images annotated by known compound mechanism achieved 85% accuracy in MoA classification. Notably, misclassification patterns showed convergence among compounds with related mechanisms, indicating that the model captured biologically meaningful phenotypic relationships rather than superficial image features. By integrating controlled single-cell imaging with deep representation learning, this approach provides a robust and scalable framework for image-based MoA classification and for systematic comparison of phenotypic responses across compounds.

## Results

### Experimental Approach

To enable high-throughput MoA classification from single cell imaging, we employed a nanowell-in-microwell platform that we described previously^20–23^. Briefly, nanowells are formed on a poly-d-lysine coated glass substrate by photolithographic patterning. This process involves coating UV-curable prepolymer on a rectangular glass slide, followed by UV exposure through a photomask to create nanowells with dimensions 25×25 µm. A plastic frame conforming to ANSI-standard 384-well plate dimensions is then adhered to each glass slide, where each microwell contains an array of 6,400 nanowells (80×80 grid). We performed MoA analysis on MDA-MB-231 breast cancer cells, which provide a consistent cellular background and are relatively resistant to anoikis, allowing them to be evaluated as isolated single cells. These cells were stained to highlight nuclear morphology (Hoechst 33342) and mitochondrial dynamics (MitoTracker Green FM) and seeded into the microwells (**Figure 1a**). Individual cells were confined within nanowells inside of microwells and incubated with cancer chemotherapeutic compounds of known MoA. A collection of 21 compounds spanning 11 MoA classes were included in this analysis. (**Figure 1b**, **Table 1**). The nanowell array in each microwell was imaged by automated high-resolution fluorescence microscopy (**Figure 1c**). Each microscopy field is then segmented into individual nanowell images to produce >500,000 single-cell images to train and validate a ResNet-18 CNN (**Figure 1 d-e**).

**Figure 1.**
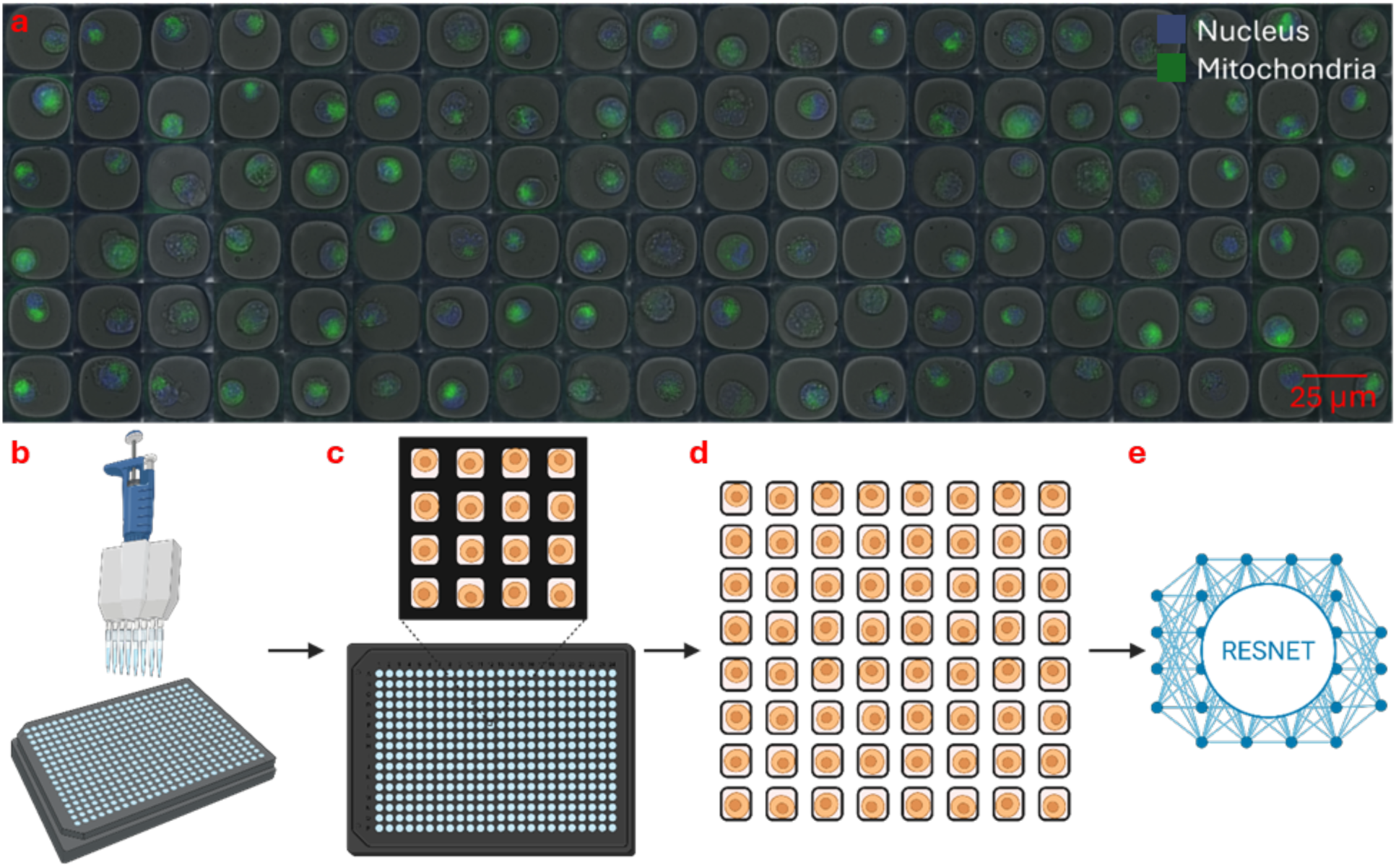
Single-cell Drug Screening. **(A)** Example images of MDA-MB-231 cells deposited in 25x25 µm glass-bottom nanowells. Cells have been fluorescently stained to highlight the nucleus (blue) and mitochondria (green). Cells in each column has been treated with a different drug, with the final column representing untreated controls. **(B-E)** Experimental process outlining dispensing cells using a standard multi-channel pipette with the ANSI conforming nanowell plates. Post imaging, nanowells are segmented into a database of single-cell images which are then analyzed using a ResNet-18 model.

**Table 1.** Reference compounds used in this study with established mechanisms of action. ^24^.

| **Ref** | **Name** | **Primary Mechanism of Action** | **Off-Target Effects** | **Reported IC<sub>50</sub> (µM)** |
|---|---|---|---|---|
| **A1** | Perifosine | AKT | MAPK/ERK modulation, JNK & p38 activation, PKC effects, lipid raft disruption | 0.6-8.9<br>[Source] |
| **A2** | GSK690693 | AKT | AGC kinases (PKA, PKC), CAMK family (AMPK, DAPK3), STE kinases (PAKs) | 0.002-0.081<br>[Source] |
| **B1** | Crizotinib | ALK, c-MET, ROS1 | TGFβR1, JAK/STAT, MAPK | 0.79-10.8<br>[Source] |
| **B2** | Lorlatinib | ALK, ROS1 | Can inhibit other kinases | 0.014-0.113<br>[Source] |
| **C1** | Palbociclib | CDK4/6 | ERK5, JNK, CDK16/17, PI3KC3 | 0.58-14.6<br>[Source] |
| **C2** | K03861 | CDK2 | CDK8, CDK19 | 0.0097-0.134<br>[Source] |
| **D1** | Gefitinib | EGFR | HER2 (weak) | 4.1-45.31<br>[Source] |
| **D2** | Erlotinib | EGFR | HER2 (weak) | 0.4-55.51<br>[Source] |
| **E1** | PD0325901 | MEK | -- | 0.00033-0.001<br>[Source] |
| **E2** | U0126 | MEK | -- | 0.06-0.07<br>[Source] |
| **F1** | Rapamycin | mTORC1 | mTORC2 | 0.9-35.79<br>[Source] |
| **F2** | Everolimus | mTORC1 | mTORC2 | 0.005-0.006<br>[Source] |
| **G1** | Sunitinib | Mutli-kinase | Broad kinase inhibition | 2.6-32<br>[Source] |
| **G2** | Sorafenib | Mutli-kinase | Broad kinase inhibition | 0.94-36<br>[Source] |
| **H1** | Wortmannin | PI3K | mTOR, DNA-PK, MLCK, MAPK | 0.003-1.8<br>[Source] |
| **H2** | LY294002 | PI3K | PIM1 kinase, mTOR | 0.38-9.7<br>[Source] |
| **I1** | Marizomib | Proteasome | -- | 0.0035-0.430<br>[Source] |
| **I2** | Bortezomib | Proteasome | -- | 0.003-16.65<br>[Source] |
| **J1** | Galunisertib | TGF-β | -- | 0.056<br>[Source] |
| **K1** | Dasatinib | Tyrosine kinase | Broad kinase inhibition | 0.112-12<br>[Source] |
| **K2** | Imatinib | Tyrosine kinase | Broad kinase inhibition | 37<br>[Source] |

### Generation of annotated training dataset using drugs of known mode of action

To train deep learning models of MoA classification, we stained MDA-MB-231 cells using Hoechst 33342 and MitoTracker Green FM to highlight the nucleus and mitochondria before seeding the cells into nanowell-in-microwell plates. Cell samples treated with each of the 21 drugs spanning 11 previously defined MoAs (**Table 1**), across eight concentrations spanning 0.01 nM to 100 μM. The treated cells were imaged at 1h, 4h, 8h and 12h post-treatment. Collectively, we collected >600,000 images of single nanowells with individual cells. The raw images were then filtered to remove empty wells, clumped cells, and low signal images. Following these pre-processing steps, a dataset of 506,853 single-cell images, annotated with reference drug MoA, was compiled to train the deep learning model.

### Training of deep learning model for classification of mechanism of action

To evaluate whether a machine learning model could classify compounds based on MoA, we annotated cell images with the 11 established MoA of the treated compound to trained a convolutional neural network (CNN) based on ResNet-18^25^ for MoA classification. ResNet-18 is an 18-layer deep convolutional network with residual connections between layers. Four separate models were trained using images acquired at 1h (T01), 4h (T04), 8h (T08), and 12h (T12) post-treatment. (**Figure 3a**). The models were trained over 10 epochs, wherein training accuracy for all treatment time points exceeded 94% (**Figure 3a**). For each of the compounds screened, 20% of the cell images were held out from the training dataset and used for validation. These validation samples achieved classification accuracies of 89% for T01, 85% for T04, 81% for T08, and 83% for T12 (**Figure 3a**). Classification performance and the mechanism-aligned misclassification trends were consistent across all four timepoints, indicating that the main conclusions do not depend on the choice of timepoint. We focused on T04 as a representative intermediate timepoint because it allows drug-induced phenotypes to develop while limiting the progressive loss of viable cells seen at later timepoints (the number of usable single-cell images declined from 180,188 at T01 to 65,877 at T12 as dead cells were removed from the well plate). We built a confusion matrix to show how the model misclassifications behave (**Figure 3b**). The T04 model showed a range of MoA class accuracies ranging from 68-94%. Interestingly, misclassifications were not randomly distributed but followed very specific trends. We expected some classification overlap between untreated and treated groups, as low-dose conditions may induce only subtle changes in cellular phenotypic signatures. However, the misclassification between classes seems to be related to overlapping molecular pathways. Overall, the model trained on data from T04 achieved an average precision of 86%, recall of 85% and F1-score of 86% (**Table 2**). We further explored the structure of the learned feature space of this CNN by extracting embeddings from the hidden layer preceding the final output layer of the CNN and visualizing them in a t-SNE (**Figure 4**). Together, these validation experiments indicate that CNN models can reliably suggest the MoA of drugs directly from images of treated cells isolated in nanowells.

**Figure 2.**
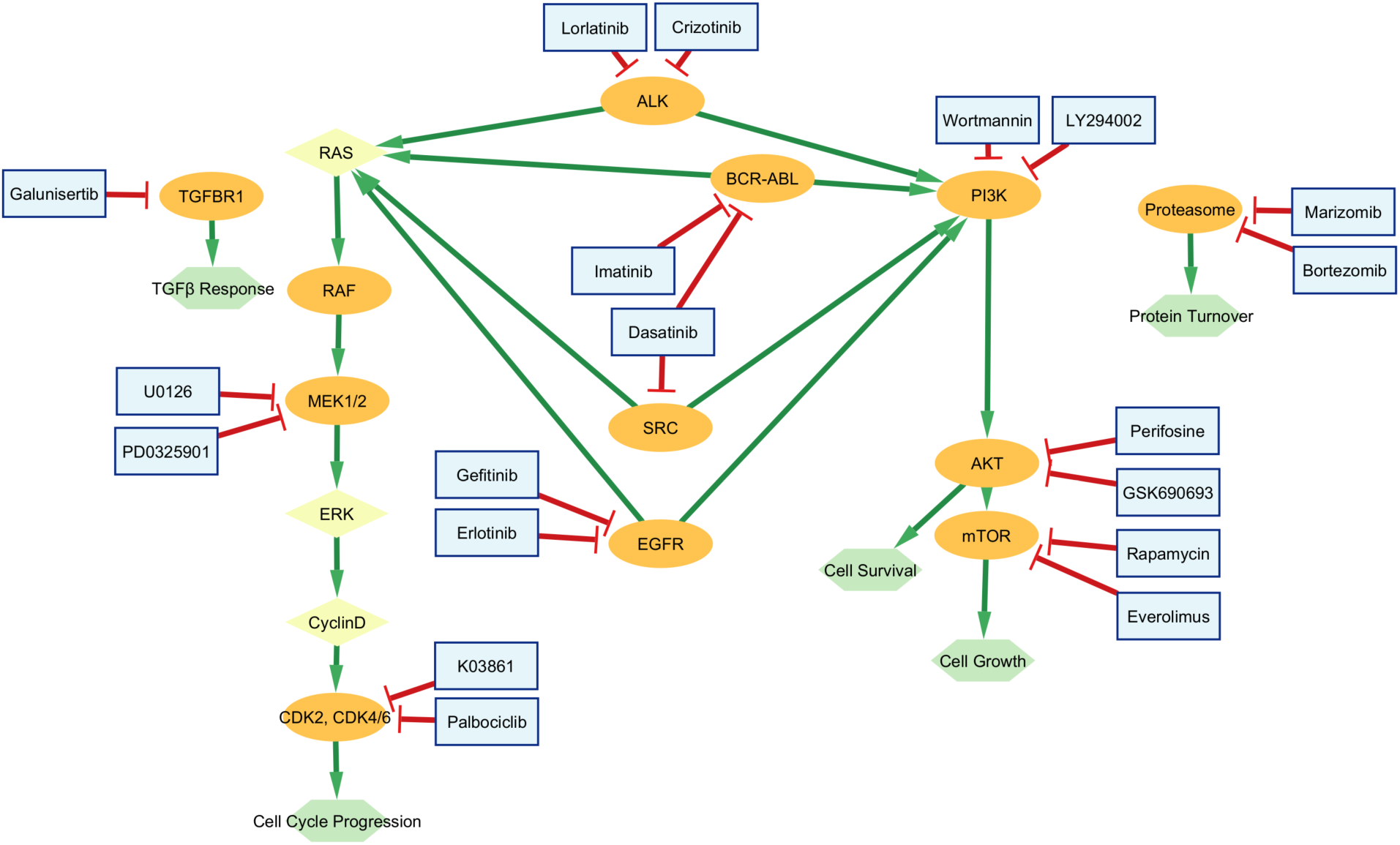
Drug Targets. Schematic overview of the oncogenic signaling pathways targeted by the pharmacological inhibitors used in this study.

**Figure 3.**
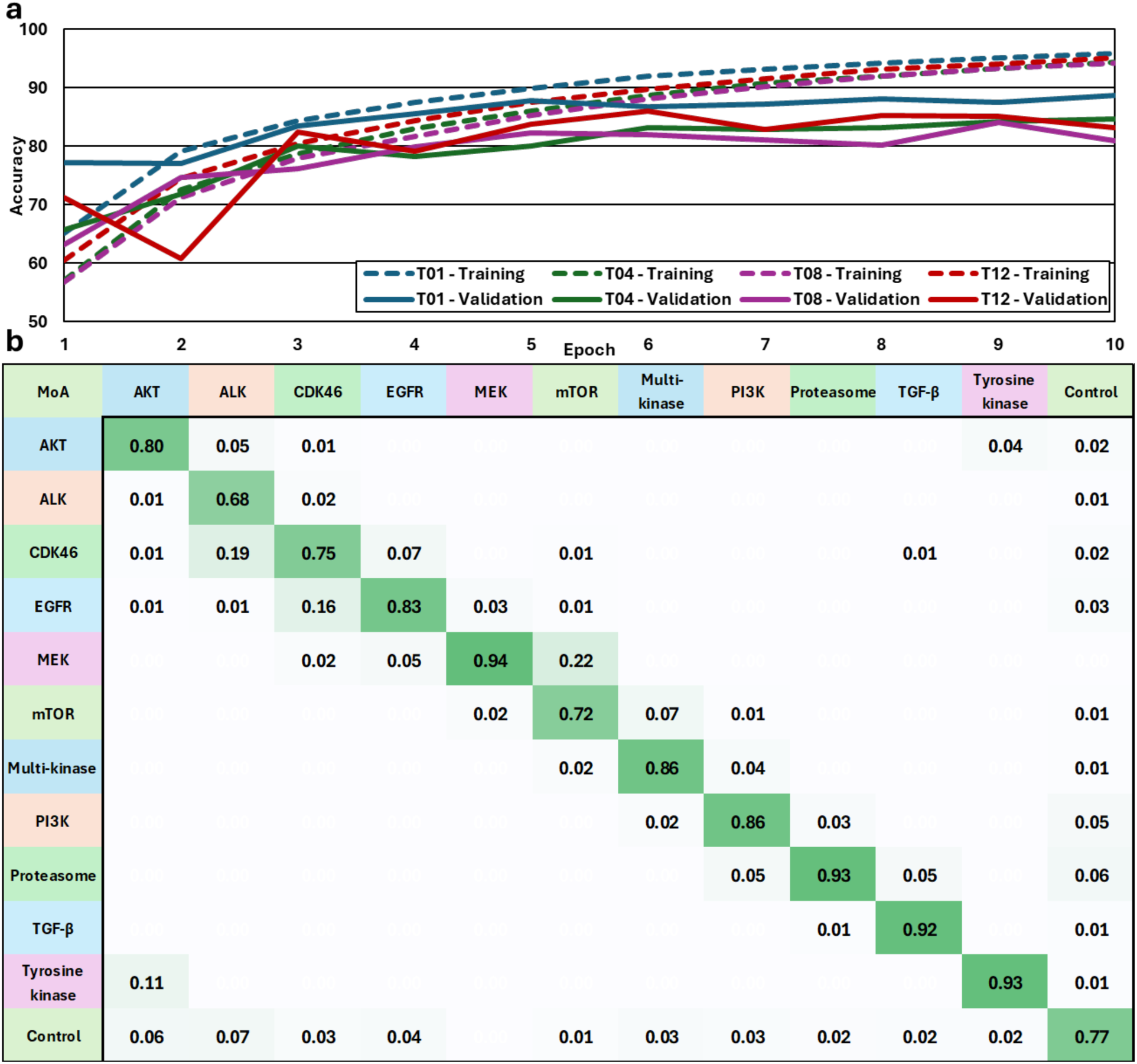
Supervised Learning for Mechanisms of Action. MDA-MB-231 were treated with 21 reference compounds and images were acquired of nanowells with single treated cells. A ResNet-18 model was trained with cell images that were annotated for the known mechanism of action for each compound. **(A)** MoA training and validation classification accuracy of cells imaged at 1h (T01), 4h (T04), 8h (T08), and 12h (T12). **(B)** Confusion matrix between classes. Entries <0.005 were left empty.

**Figure 4.**
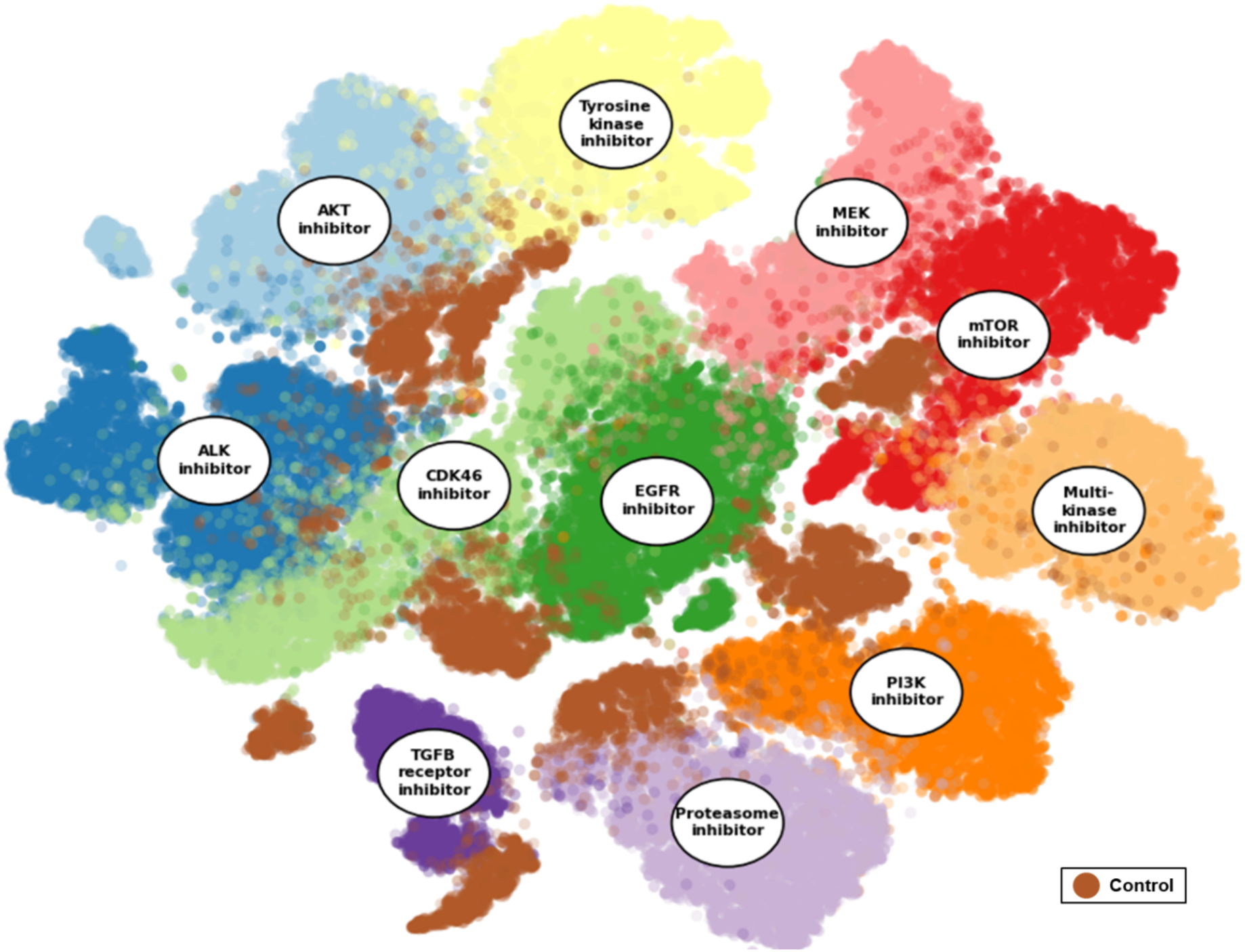
Supervised learning separates drug mechanisms of action in phenotypic feature space. t-SNE projections of deep image features learned by the MoA classifier show distinct clusters corresponding to drug classes. Each point represents a single cell, colored by MoA, with class labels placed at cluster centroids. Related inhibitor classes form neighboring clusters, while controls were distinct but broadly distributed, demonstrating that the model captures MoA-specific phenotypic signatures from non-specifically stained images.

**Table 2.** MoA classification performance for models trained on established drug MoA.

| **Class** | **Precision** | **Recall** | **F1-Score** |
|---|---|---|---|
| **AKT inhibitor** | 0.83 | 0.92 | 0.87 |
| **ALK inhibitor** | 0.83 | 0.88 | 0.85 |
| **CDK46 inhibitor** | 0.68 | 0.85 | 0.75 |
| **EGFR inhibitor** | 0.82 | 0.74 | 0.78 |
| **MEK inhibitor** | 0.89 | 0.83 | 0.86 |
| **mTOR inhibitor** | 0.84 | 0.88 | 0.86 |
| **Multi-kinase inhibitor** | 0.92 | 0.85 | 0.89 |
| **PI3K inhibitor** | 0.89 | 0.92 | 0.90 |
| **Proteasome inhibitor** | 0.89 | 0.93 | 0.91 |
| **TGFB receptor inhibitor** | 0.97 | 0.89 | 0.93 |
| **Tyrosine kinase inhibitor** | 0.95 | 0.85 | 0.90 |
| **Control** | 0.85 | 0.69 | 0.76 |
| **Average** | **0.86** | **0.85** | **0.86** |

### Convergent morphological phenotypes among drugs of common MoAs

To evaluate whether our deep learning strategy would correctly classify the effect of drug treatment without prior knowledge about MoA, we trained a second ResNet-18-based CNN to classify cell images annotated by 21 compound identities rather than MoA. We hypothesized that drugs of similar MoA might induce convergent morphological phenotypes within single cells and that misclassified compounds would more frequently identify the drug with similar MoA. We trained a 21 class ResNet-18 model using the same cell images annotated with compound identity instead of MoA. We once again trained four models over 10 epochs using data from T01, T04, T08 and T12. The models achieved training accuracy ranging between 93-95% and validation accuracy between 81-88% (**Figure 5A**). Specifically, at timepoint T04, the model achieved an average precision of 72%, recall of 71% and F1-score of 71% (**Table 3**). These metrics are lower than when the model is trained solely on MoA classes, which is expected given the increased likelihood of misclassification among compounds sharing a common MoA. Interestingly, grouping the predictions based on MoA improved model performance, resulting a 6% increase in accuracy to achieve an average validation accuracy of 87% (**Figure 5b**).

**Figure 5.**
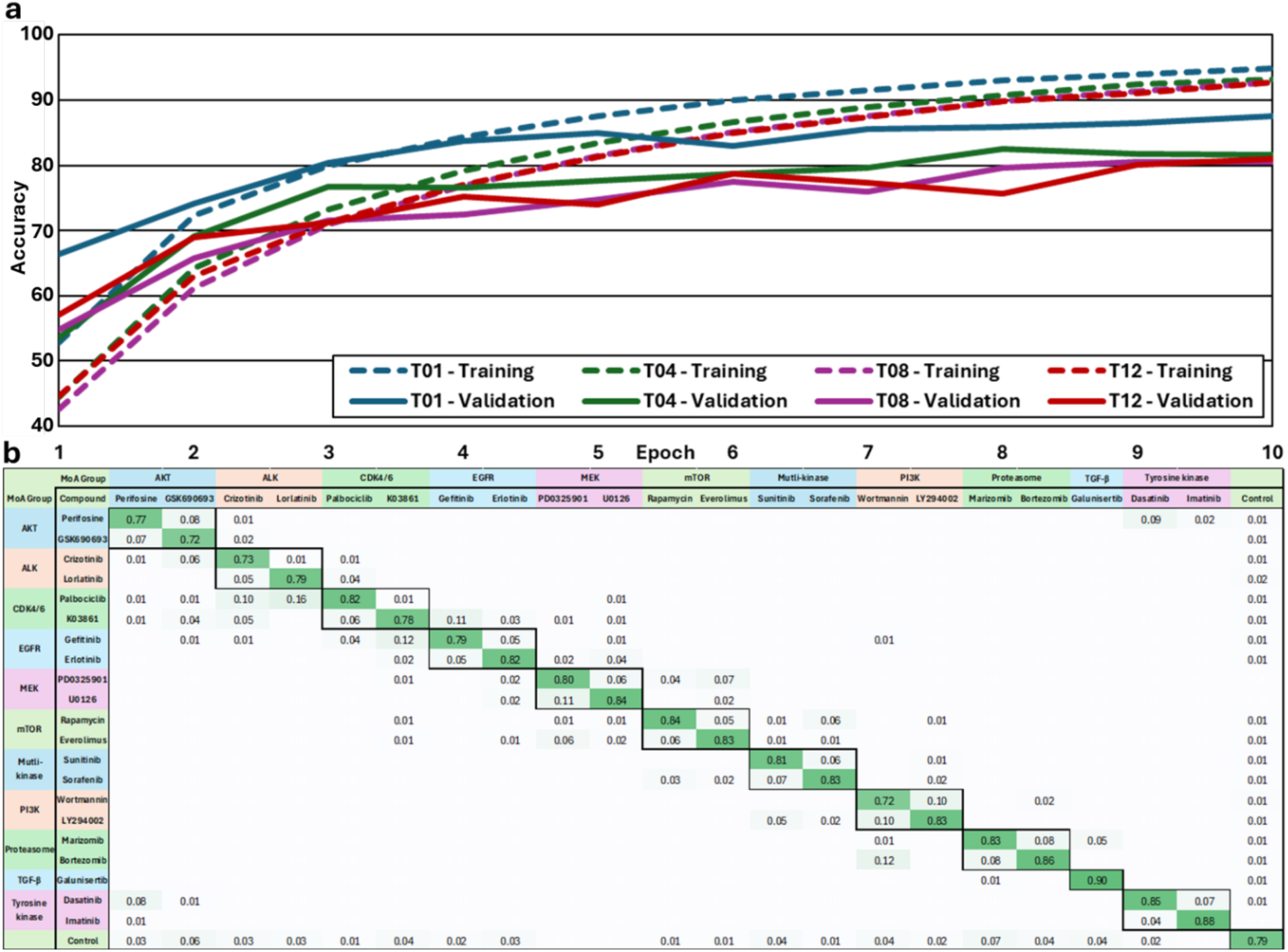
Supervised Learning on Overlapping Compounds. MDA-MB-231 were treated with 21 reference compounds and images were acquired of nanowells with single treated cells. A ResNet-18 model was trained with cell images that were annotated for drug identity. **(A)** Training and validation curves for models trained on data from each imaging timepoint. **(B)** Confusion matrix indicating the accuracy of drug classification as well as the frequency that drug misclassification instead identified a drug with the same mechanism of action. Table entries which were <0.005 where left empty.

**Table 3.** MoA classification performance for models trained on annotated drug identity.

| **Class** | **Precision** | **Recall** | **F1-score** |
|---|---|---|---|
| Perifosine | 0.77 | 0.78 | 0.78 |
| GSK690693 | 0.72 | 0.85 | 0.78 |
| Crizotinib | 0.73 | 0.89 | 0.80 |
| Lorlatinib | 0.79 | 0.86 | 0.82 |
| Palbociclib | 0.82 | 0.68 | 0.75 |
| K03861 | 0.78 | 0.64 | 0.71 |
| Gefitinib | 0.79 | 0.73 | 0.76 |
| Erlotinib | 0.82 | 0.81 | 0.82 |
| PD0325901 | 0.80 | 0.78 | 0.79 |
| U0126 | 0.84 | 0.83 | 0.83 |
| Rapamycin | 0.00 | 0.00 | 0.00 |
| Everolimus | 0.84 | 0.82 | 0.83 |
| Sunitinib | 0.72 | 0.72 | 0.72 |
| Sorafenib | 0.72 | 0.72 | 0.72 |
| Wortmannin | 0.72 | 0.71 | 0.71 |
| LY294002 | 0.72 | 0.69 | 0.70 |
| Marizomib | 0.71 | 0.68 | 0.69 |
| Bortezomib | 0.71 | 0.68 | 0.69 |
| Galunisertib | 0.70 | 0.68 | 0.69 |
| Dasatinib | 0.69 | 0.68 | 0.68 |
| Imatinib | 0.68 | 0.66 | 0.67 |
| Control | 0.67 | 0.65 | 0.66 |
| **Average** | **0.72** | **0.71** | **0.71** |

We further explored the structure of the learned feature space of this CNN by extracting embeddings from the hidden layer preceding the final output layer of the CNN and visualizing them in a t-SNE (**Figure 6**). We illustrated each pair of drugs with shared mechanism by drawing a line between cluster labels. We observed that drugs with similar MoA frequently emerged in proximity to one another in the CNN learned feature space, indicating that they induce morphologically similar changes in single cells. For example, tyrosine kinase inhibitors (TKIs) sunitinib and sorafenib cluster tightly and both compounds have shared mechanisms of VEGF and PDGF receptor tyrosine kinase inhibition^26^. In contrast, the MOA-paired compounds lorlatinib and crizotinib both target ALK and ROS1 but formed more distant clusters, which may reflect the pleiotropic effects of lorlatinib that selectively influence cell morphology, adhesion and motility^27^.

**Figure 6.**
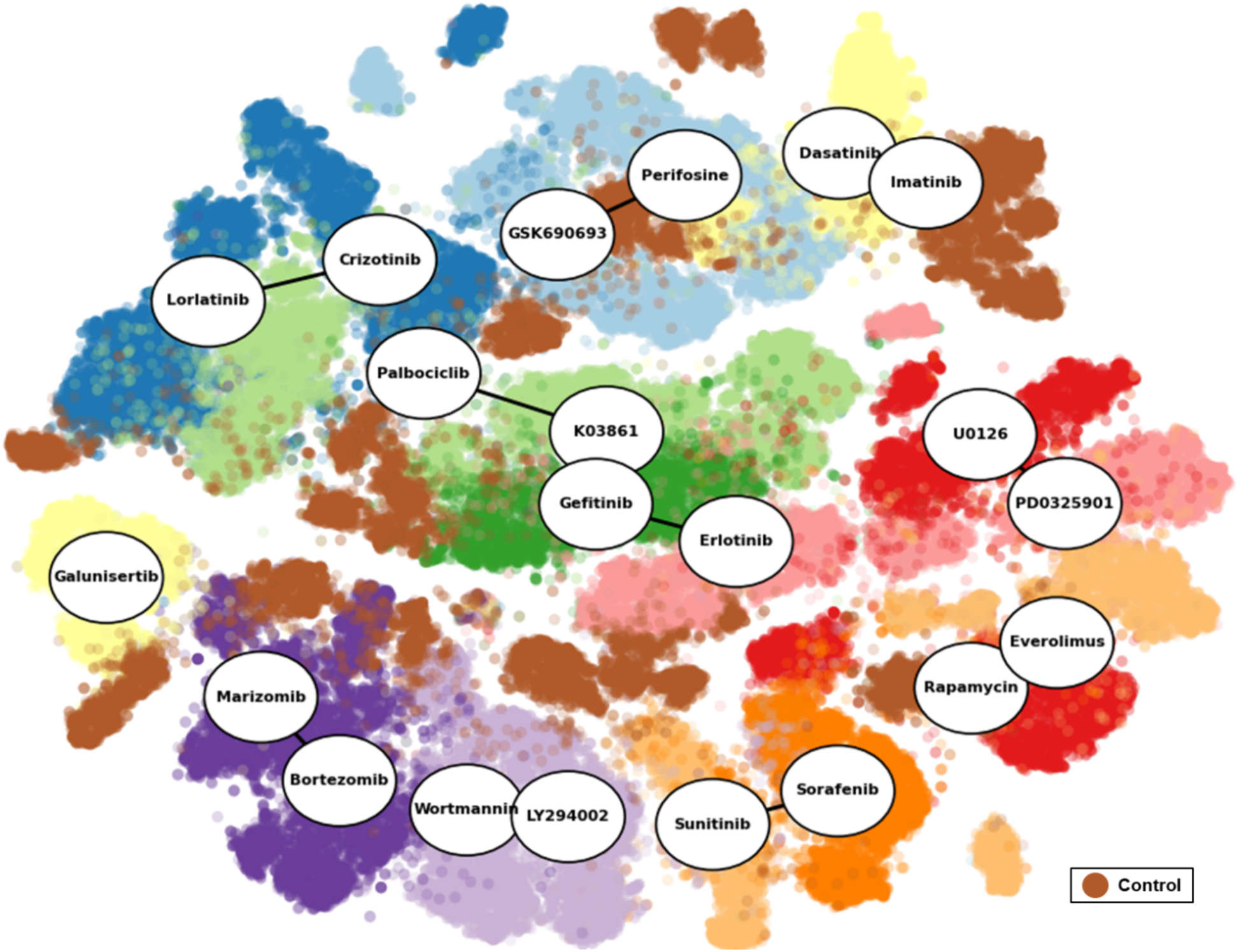
Supervised Learning on Overlapping Compounds. t-SNE embeddings projected onto a 2D space with the drug treatment for each class centered on the centroid of the data. A line has been drawn between compounds with the same MoA.

Crizotinib and lorlatinib were commonly misclassified as palbociclib (10% and 16% respectively), which is shown in **Figure 5b**. However, this misclassification is biologically consistent since ALK signaling lies upstream of CDK4/6, and blocking ALK often suppresses CDK4/6 activity, resulting in cell-cycle arrest^28^. Interestingly, the ALK inhibitors that have upstream targets 6.2 times more likely to be misclassified as CDK4/6 inhibitors than the reverse, which suggests that confusion with upstream targets may be less frequent than downstream targets in the same pathway.

### Investigating time and dose dependent contributions

To evaluate the effects of drug dose and incubation time on CNN performance, we trained 30 separate ResNet-18 models using the same dataset as the previous model, but with cell images stratified by both dose and incubation time. Models were trained independently for each time point and concentration. Across timepoints T01, T04, T08, and T12, the dataset contained 180,188, 142,116, 126,722, and 65,877 images, respectively. Dataset size decreased over time as increasing cell death led to exclusion of morphologically degraded cells, reducing available training data and contributing to poorer model performance despite stronger treatment effects. Although higher drug concentrations and longer exposure times were expected to improve classification accuracy by amplifying phenotypic differences, they also led to an increased prevalence of dead cells. The training and validation accuracies of each model were assessed after 10 epochs, where training accuracy ranged from 89.5% to 96.6% and validation accuracy ranged from 68.5% to 90.3%, respectively (**Figure S1**).

To further explore how single-cell morphology varied across conditions, t-SNE projections were generated using features extracted from the trained models (**Figure 7**). At lower concentrations and shorter exposure times, morphological profiles from different treatments exhibited considerable overlap, reflecting difficulty in discerning between subtle phenotypic changes. As drug concentration and exposure time increased, clusters corresponding to different treatments became progressively more distinct and separated. This pattern indicates that drug-induced phenotypic divergence is both dose-and time-dependent, with higher doses and longer treatments amplifying cellular responses to perturbation. Notably, the biologically relevant clustering was most often preserved with moderate treatments over longer periods of time.

**Figure 7.**
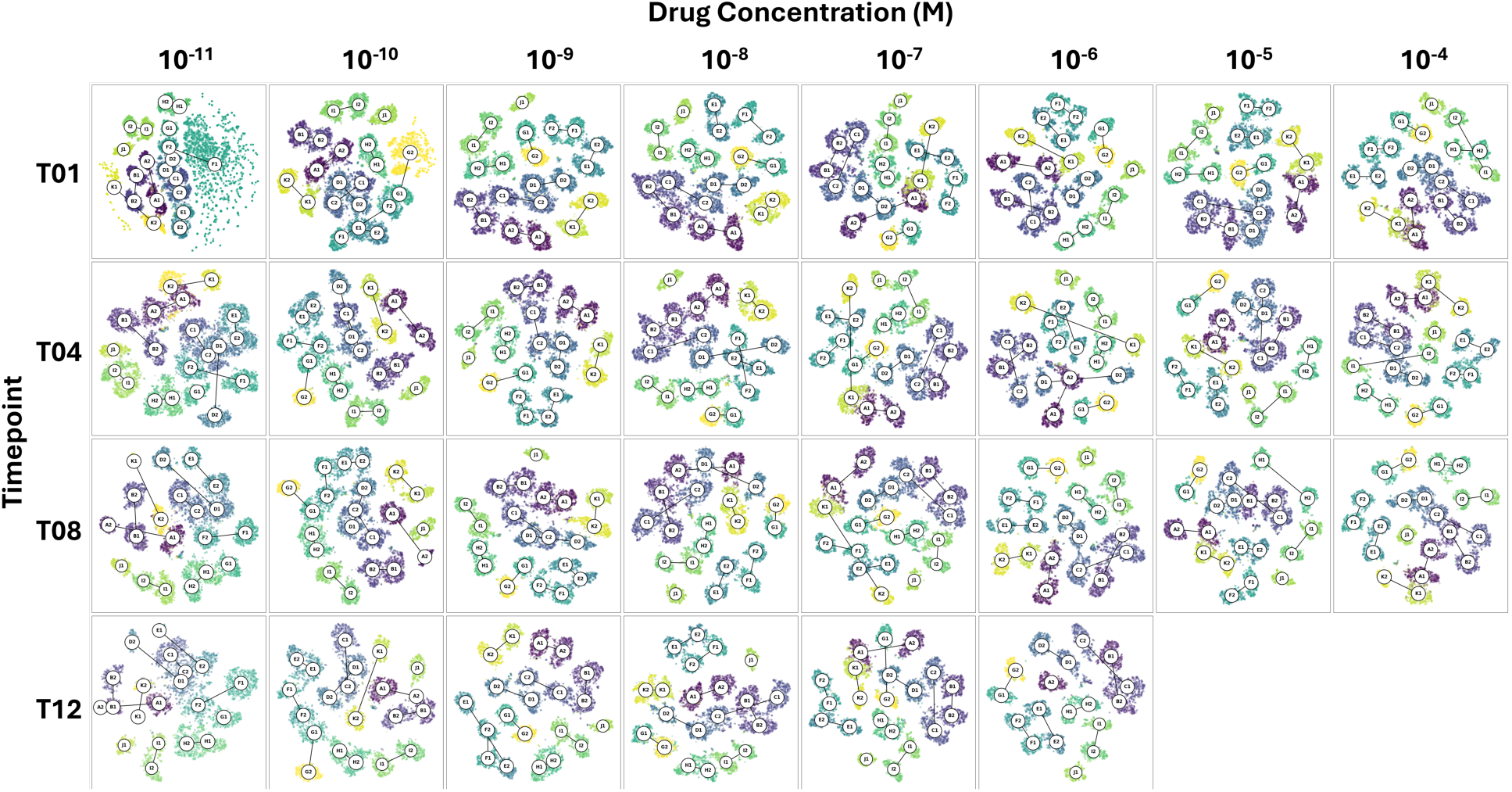
Condition Clustering. t-SNE projections of single-cell morphological data across varying drug concentrations and timepoints with clusters labeled according to Table 1. Each panel corresponds to a specific condition, with colors representing different drug treatments. At higher doses, clusters become more defined and separate, indicating increased phenotypic divergence in response to treatment.

## Discussion

Here, we described a deep learning strategy for mechanism-of-action discovery enabled by isolating cells within nanowells. Our imaging platform consists of an array of nanowells fabricated inside each microwell of standard 384-well glass-bottom microwell plates, which are compatible with available microscopy equipment. Confining single cells in nanowells enables standardized imaging of each cell and eliminates the need for cell segmentation, which is error-prone and constitutes a primary bottleneck to image-based MoA discovery. Using this platform, we generated >500,000 annotated single-cell images across 21 cancer chemotherapeutic compounds spanning 11 known MoA, exposed to 30 combinations of doses and incubation times. We showed that CNNs trained on these images could classify MoA with 85% accuracy.

A key advantage of the nanowell-based workflow for prediction of MoA is its capability for high-throughput single cell analysis without needing cell segmentation and feature extraction. In contrast to conventional high-throughput MoA prediction workflows that quantify aggregate features of a cell population in standard microwells^29^, training CNN models based on single cell images ensures that the heterogeneity of the cell population is integrated into the analysis. Other deep learning models that are trained on single cell images acquired their training data by segmentation of single cell images from pre-existing image databases^30^ or from high-throughput imaging. However, single cell segmentation is severely complicated by cell overlap and imaging artifacts, to the extent it creates a bottleneck in the analytic workflow and injects noise into downstream analysis. These workflows then rely on feature extraction to derive hundreds to thousands of predefined cell measurements, and although classification models built on such features have shown encouraging results, their accuracy depends heavily on feature selection and extraction quality^31^. By imaging single cells in nanowells, our platform bypasses the need for cell segmentation by training on the entire nanowell image, and it eliminates cell-density bias by isolating one cell per well to standardize cell density. A limitation of our platform is that nanowell contact may alter cell morphology but we observed this effect in <10% of cells and did not see evidence that these effects impaired cell classification. In place of predefined endpoints such as nuclear area or mitochondrial intensity, our AI model performed pixel-level classification, learning feature representations directly from the raw images, so that classification is driven by learned patterns of intensity, texture, and spatial organization rather than human-defined measurements. Therefore, isolating single cells in nanowells produces consistent, regularized framing for cell imaging to focus training of the AI model on biologically relevant features specific to the MoA.

Our results demonstrate that our deep learning models generated from nanowell images can accurately classify MoA of structurally diverse compounds, even though cellular response to these compounds may exhibit significant phenotypic overlap. A supervised model, where the training images were labeled with compound MoA, was able to correctly classify hold-out images with 85% accuracy. Cell images with shared MoA clustered based on shared feature space. Moreover, compounds that target distinct MoAs but share converging cell signaling pathways clustered together. For example, compounds that inhibit EGFR and MEK pathways are clustered closely, which is consistent with the phenotypic convergence expected based on their shared use of the MAPK/ERK pathway^32^. Similarly, numerous drugs targeted the pathway that is shared by PI3K/AKT/mTOR^33^. Even more significant than compound classification using MoA label is that the compounds still clustered when the model is trained on images that were not labeled directly with a MoA but only on compound identity. Compounds were identified with 81% accuracy, but it is noteworthy that compounds that were incorrectly identified were typically mis-identified as the compound with a shared target. This finding suggests that it may be possible to infer the MoA of unknown compounds based on the association with known compounds to which they are classified. We note that image-based MoA analysis is intended to classify compounds against previously characterized mechanisms rather than to discover new ones. Within this scope, a key result is that the model recovers mechanism-level organization without prior knowledge of MoA. In fact, when a second AI model was trained only on compound identity, its errors and learned embeddings group compounds by shared mechanism, indicating that regularized single-cell morphology encodes MoA structure that need not be specified in advance.

An important consideration in image-based profiling is the potential for batch effects, such as illumination gradients or well-position effects, which can lead models to classify on technical rather than biological features^17^. Several observations indicate that classification in this study is driven by compound biology rather than technical signal. Compounds that share a mechanism of action occupy different regions of the plate, yet they converge both in the model trained on MoA labels and in the model trained on compound identity alone. In the compound-identity model, misclassifications occur preferentially between compounds that share a mechanism rather than between compounds that are physically adjacent on the plate. If illumination or well-position effects were driving the models, errors and clustering would be expected to track plate location; instead, they track mechanism of action, a pattern difficult to explain by batch effects. Consistent with this interpretation, misclassified control cells are distributed approximately evenly across many compound classes rather than being systematically confused with a single neighboring condition, as a localized positional artifact would produce. This even distribution instead reflects the heterogeneity of the untreated baseline phenotype, which overlaps partially with the many low-concentration conditions that produce only subtle changes in cell state. A further consideration is that confining cells as isolated single cells is not equally compatible with all cell populations. Cell types that depend on cell-cell contact or attachment to extracellular matrix for survival may undergo anoikis when maintained as isolated single cells, which would limit the applicability of the current configuration to such populations.

Our approach addresses key limitations in existing morphological profiling strategies by integrating single-cell nanowell imaging with deep learning-based classification. Traditional methods, such as the CellProfiler pipeline^34^, are constrained by reliability of cell segmentation, as well as the reliance on human-engineered features, which may be limited by human cognition and description. By isolating individual cells in regularly spaced nanowells, we enable CNNs to be trained directly on single-cell images without requiring cell segmentation. This approach avoids errors that commonly arise when analyzing overlapping or clustered cells. Our nanowell-based machine learning framework for MoA prediction supports a wide range of applications that combine scalable imaging with deep learning at single-cell resolution. Future directions include expanding the compound library to encompass agents with subtle or mixed phenotypic signatures and applying the framework to diverse cell types to test generalizability across biological contexts. Moreover, incorporating time-lapse imaging could capture dynamic cellular responses and early phenotypic changes. Ultimately, this approach enables a more systematic exploration of drug-induced phenotypes, facilitates novel target identification, and contributes to safer and more effective therapeutic strategies.

## Methods

### Cell Culture

MDA-MB-231 (ATCC HTB-26) breast cancer cells were used in this study. Cells were cultured in Dulbecco’s Modified Eagle Medium (DMEM) with 4.5 g/L d-Glucose and l-Glutamine (Gibco), supplemented with 10% Fetal Bovine Serum (FBS, Gibco) and 1X Penicillin/Streptomycin (P/S, Gibco). Cells were incubated in a T-75 flask (Corning) at 37 °C with 5% CO2 in a 90% humidified incubator. When needed, the adherent cell lines were released from the flasks with Trypsin-EDTA (0.25%, Gibco) and then washed twice in complete media, prior to use.

### Sample Preparation

The library of compounds was purchased from Millipore sigma and includes: Perifosine (SML0612), GSK690693 (SML0428), Crizotinib (PZ0191), Lorlatinib (PZ0039), Palbociclib (PZ0383), K03861 (SML2971), Gefitinib (SML1657), Erlotinib (SML3621), PD0325901 (PZ0162), U0126 (662005), Rapamycin (553210), Everolimus (SML2282), Sunitinib, (PZ0012), Sorafenib (SML2653), Wortmannin (681675), LY294002 (440202), Marizomib (SML1916), Bortezomib (5.04314), Galunisertib (SML2851) Dasatinib, (SML2589), and Imatinib (CDS022173). Sample dilutions were prepared in u-bottom 96-well plates prior to the experiment. 10 µL of the compound dilutions were initially pipetted into the working nanowell plates. Next MDA-MB-231 cells were resuspended in a falcon tube containing DMEM media, Hoechst 33342 (H3570, Invitrogen) and MitoTracker Green FM (M7514 Invitrogen). 70 µL of the cell containing solution were dispended into each well of the 384 well plate resulting in a concentration of 0.16 μM Hoechst 33342, 0.1 µM MitoTracker Green FM and 0.1% DMSO. Plates were immediately spun down at 100 g for 1 minute before being transferred to a Nikon Ti2 for imaging.

### Nanowell-in-Microwell Plate Design

Cells were seeded into nanowells in microwell at a density of 25,000 cells/mL and settled for 5 minutes before centrifugation at 50 × *g* for 60 seconds. To avoid edge illumination effects, compounds were not added to the most peripheral rows and columns of the plate. Compounds were added to each column in the order listed in Table 1. A 10-fold dilution series was generated in each row of the plate, where the final row was the negative control.

### Microscopy

Microscopy imaging was performed using a Nikon Ti-2E inverted fluorescence microscope. Images were acquired using a Nikon CFI Plain Fluor 10× objective and a 14-bit Nikon DS-Qi2 CMOS camera. Images were captured using three channels: brightfield with phase contrast, DAPI (Nikon C-FLL LFOV, 392/23 nm excitation, 447/60 nm emission and 409 nm dichroic mirror), and EGFP (Nikon C-FLL LFOV, 466/40 nm excitation, 525/50 nm emission and 495 nm dichroic mirror). Illumination for brightfield imaging was performed using the built in Ti-2E LED. Epifluorescence excitation was performed using a 130 W mercury lamp (Nikon C-HGFI). Gain, exposure and vertical offset were automatically determined using built-in NIS functions to avoid user biasing. Cells were imaged in 384 well format, 25×25 µm glass bottom nanowell arrays provided by ImageCyte Technologies. An automated procedure was run on NIS using the JOBS module to take 25 images spanning the entire well of each well, in each of the 3 channels. The images were exported from NIS to standard TIFF.

### Data Analysis Software

Image analysis was done in custom python scripts. The analysis started by segmenting the individual nanowells from the images using a thresholding value with was automatically obtained using the Otsu method. After thresholding the square objects were detected and the coordinates extracted for further assessment. The individual nanowell images were then analyzed to remove nanowell containing no cells, multiple cells or dead cells. Cells contacting the nanowell boundary were retained as edge contact is a regular and reproducible feature of the nanowell geometry that occurs across all treatment conditions. Excluding these cells would remove valid single-cell observations and could introduce selection bias. The library of single-cell images was then saved in a lossless format for machine learning training.

### AI Model Training

Each network was trained for 10 epochs using the Adam optimizer with a learning rate of 0.001 and a batch size of 32. Categorical cross-entropy, implemented via a softmax output layer, was used as the loss function for backpropagation. Training was performed in Python (v3.10) using PyTorch (v2.6.0) on a Windows 10 workstation equipped with an Intel i5-7500 CPU (3.40 GHz), 32 GB DDR4 RAM (3200 MHz), and an NVIDIA GTX 1080 GPU with 8 GB of memory.

### Training Performance Evaluation

To identify the miss-classifications between labels, a confusion matrix was constructed. All of the images in the testing dataset were used. A class label was predicted using the CNN and compared against the true class label. The diagonal positions correspond to true labels, while the off-diagonal positions show miss-classifications. The matrix was normalized by the number of images in each class to represent probabilistic accuracy.

### Data Clustering and Visualization

To generate low-dimensional representations of the learned feature space, we applied t-distributed stochastic neighbor embedding (t-SNE) to feature vectors extracted from the trained model. Specifically, 512-dimensional feature vectors were obtained from the fully connected layer immediately preceding the final classification layer following model training. The final classification layer was removed, and all images in the test dataset were passed through the truncated network to compute feature representations. These feature vectors were then projected into two dimensions using t-SNE, a non-linear dimensionality reduction method that preserves local relationships in high-dimensional data.

## Author Contributions

HM supervised the study. SGB and HM conceived the idea. SGB and KM performed the experimental work. SGB analyzed the data. SGB, HM and SPD wrote the manuscript.

## Declaration of generative AI and AI-assisted technologies in the manuscript preparation process

During the preparation of this work the author(s) used ChatGPT to revise portions of the manuscript to improve writing clarity and flow. After using this tool, the authors reviewed and edited the content as needed and take full responsibility for the content of the published article.

## Supplemental Figures

**Figure S1.**
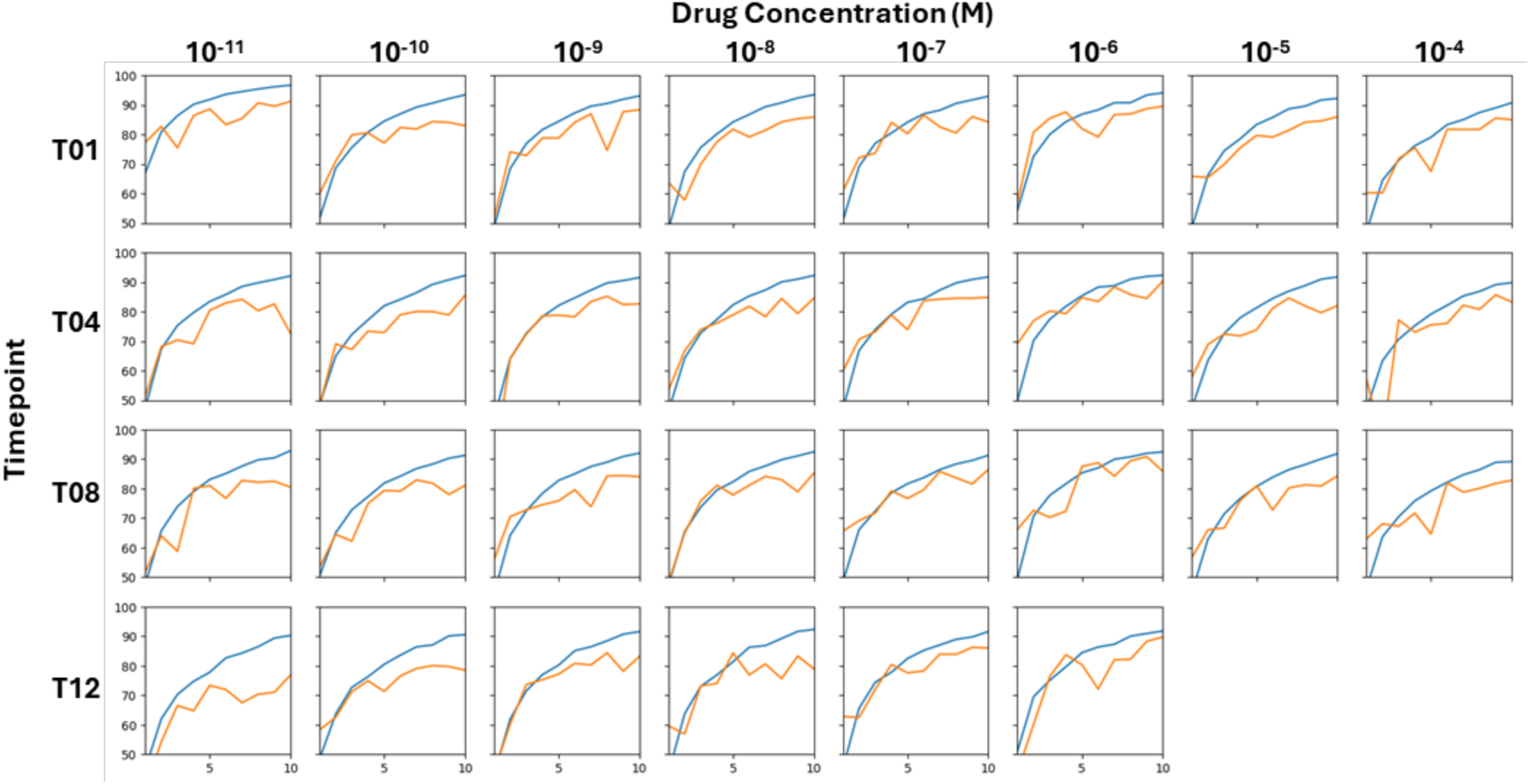
Training Results on Mechanisms-of-Action. Training and validation accuracy of ResNet18 models trained to classify single-cell images by drug treatment. Each subplot represents a separate model trained for a specific drug concentration and timepoint. Training (blue) and validation (orange) accuracies are plotted over 10 epochs.

